# Contribution Evenness: A functional redundancy metric sensitive to trait stability in microbial communities

**DOI:** 10.1101/2020.04.22.054593

**Authors:** Taylor M. Royalty, Andrew D. Steen

## Abstract

The concept of functional redundancy has received considerable attention in both the macroecology and microbial ecology literature. As a result, multiple metrics of functional redundancy have been proposed. These vary in how they weight trait levels, species abundance, functional richness, and species richness. Here we present a new functional redundancy metric tailored for community-aggregated traits, which are traits that are quantified at the community level and can be quantitatively partitioned among species. We call this metric Contribution Evenness (CE) because it measures how evenly species contribute to a community-aggregated trait. As CE is an evenness measurement, it ranges from 0 and 1, where 0 corresponds to a single species contributing to a community-aggregated trait and 1 corresponds to all species contributing equally. Using *in silico* simulations of species extinctions, we demonstrate that CE reflects the stability of an ecosystem function to species extinction, a hypothesized ecological consequence of functional redundancy. As a positive control and to illustrate how CE can be used with sequence data, we analyzed the functional redundancy of eight nitrogen-transforming pathways using 2,631 metagenome-assembled genomes from 47 marine sites. CE for marine nitrogen cycle marker genes was consistent with our qualitative understanding of which nitrogen pathways are most functionally redundant in the ocean. We found that, on average, the NH_4_^+^ assimilation pathway was the most functionally redundant (0.44 ± 0.08) while dissimilatory nitrate reduction was the least redundant (0.005 ± 0.005). As demonstrated here, CE provides a promising framework for measuring trait stability in microbiomes.

## Introduction

Recent advances in sequencing technology and sampling coverage have facilitated the discovery that important ecological functions are frequently replicated across co-occurring species in an ecosystem (1). This phenomenon is referred to as “functional redundancy”. The ecological implications of functional redundancy are not fully understood, but one hypothesized consequence is that it increases trait stability, which is the ability of microbial communities to retain a trait or function even when individual members of the community become locally extinct (2–4). To test hypotheses about the ecological and biogeochemical implications of functional redundancy in microbial communities, a quantitative and generally applicable metric is required.

Several metrics for functional redundancy exist in the literature. The diversity of metrics reflects the diverse sources of data used to calculate functional redundancy and the diverse purposes for which this concept has been applied. Functional redundancy has been measured as the ratio of community richness to the theoretical maximum number of functional entities (5), median metabolic overlap of genomic traits (6), expected variability in trait space among community members (7), the degree of covariance between functional diversity and biodiversity (e.g., Galand et al., 2018; Micheli and Halpern, 2005; Miki et al., 2014), and the simple fraction of populations in a community that bear a given function (11–13). These different metrics treat community structure differently. For instance, some metrics are sensitive to species abundances (7, 8), richness (5, 9), or neither (6). Except for the simple fraction of taxa bearing a function, the functional redundancy metrics above represent ecological function in terms of niche, where a niche is thought of as a point in a hyperdimensional trait space (14). Communities with higher taxa density in trait space are interpreted as being more functionally redundant.

This conception of functional redundancy is difficult to apply to microbial communities. There are many possible, environmentally relevant microbial functions, for most microbes these functions are known only from a microbe’s genome, and the genes for these functions are often not well-conserved. Thus, it is often impossible to define microbial niches in a meaningful and tractable way (1). As a result, existing functional redundancy metrics can be challenging to apply to microbial ecosystem.

Rather than defining functional redundancy in terms of niches, an alternative approach is to evaluate functional redundancy with respect to individual functions or traits. Here, we propose a metric designed to quantify the functional redundancy of individual traits in a microbial community, something that is currently lacking in the literature. We focus on microbial traits represented as nucleic acid sequences (DNA and RNA), i.e., gene and transcript sequences. These sequence traits are regularly measured in microbial ecology studies, are used to infer microbial metabolism, can be meaningfully compared between samples or studies, and are easily integrated into trait-based models (15–17).

To define this metric, we have treated gene and transcript sequences as “community-aggregated parameters” (18). These are parameters which are observed at the ecosystem level, but which can be quantitatively partitioned among individual species. The contribution of each species to an ecosystem-level process is calculated as the contribution of individuals to the process, weighted by their species’ abundance. Species contributions are then summed to reflect the community-level processes. Gene and transcript abundances are a natural fit for this concept, because the abundance of gene sequences can be quantified (e.g. as read depth) and assigned to specific species (e.g. by mapping reads to reference genomes or metagenome-assembled genomes) (19).

We use an evenness metric to calculate species’ relative contribution to an observed community-level parameter (20). Briefly, the “true diversity” of contributors is calculated using species contributions and traditional diversity metrics (21, 22). The observed diversity is normalized to a theoretical maximum diversity (richness) and made “unrelated”, in the sense used by Chao and Ricotta (20), to community richness. Our approach can be understood as measuring functional redundancy because it measures how evenly species in a community contribute to an ecosystem parameter—the more evenly species contribute to an ecosystem parameter, the more functionally redundant the community is with respect to that process. Accordingly, we term our measure of functional redundancy “contribution evenness” (CE).

To demonstrate the efficacy of CE, we first performed simulations to optimize the value of a constant called diversity order which is required in CE calculations (21, 22). Second, we performed an *in silico* experiment to test how well CE captures trait stability compared to other, existing functional redundancy metrics. Finally, we demonstrate the applicability of CE with real-world data by analyzing functional redundancy for eight nitrogen-transforming pathways found in metagenomic data (23) in *TARA* Oceans draft metagenome-assembled genomes (MAGs) (24). Overall, we found CE appealing because it is easily calculated from conventional shotgun genomic sequence data sets, provides insights on the relative contribution by community members to community-level processes, and is sensitive to the stability of traits to species extinction.

## Methods and Theory

### Definition of Functional Redundancy

We define a microbial community, *C*, with *S* species, where the *i*^th^ species has a relative abundance, *p*_*i*_. (For simplicity, we refer to ecological units as species, although it would be possible to perform this analysis at other taxonomic levels – strains, say, or genera – or even to perform the analysis at the level of guilds in order to evaluate resource competition among guilds, e.g., sulfate-reducers competing with methanogens for hydrogen electrons). We therefore represent *C* as:

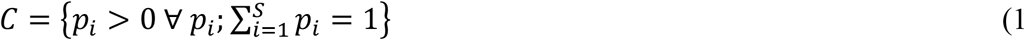

where *p* is the set of relative abundances of species, such that any given relative abundance for the *i*^th^ species is greater than zero and all species relative abundances sum to one. Equation 1 represents a traditional abundance distribution (25).

Our goal was to calculate how evenly different species in a community, *C* (equation 1), contribute to a community-aggregated parameter, or ecosystem function, *T*, at a fixed time and space. To do this, we first determined the relative contribution, *τ*_*i*_, that the *i*^th^ species makes towards *T* such that:

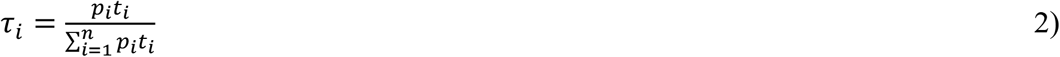

where *t*_i_ is the species-specific contribution level and *p*_i_ is the relative abundance the *i*^th^ species in *C*. In equation 2, *τ*_i_ represents the relative contribution that the *i*^th^ species contributes to a community-aggregated parameter. We note that the functional analysis and preprocessing of metagenomes and transcriptomes varies in the literature and public databases. For example, the *TARA Oceans* metagenomes have been analyzed as assembled, un-binned metagenomes, where individual genes were assigned taxonomy, annotation, and abundances (26), and as metagenome-assembled genomes (24, 27). Although the fundamental meaning of *τ* does not change across these different datasets, the *in silico* approach to calculating *τ* may slightly differ depending on how sequence data is preprocessed. For un-binned metagenomes, *τ*_*i*_ simply corresponds to the proportion of reads associated with the *i*^th^ species which map to function, *T*. Similarly for transcriptomes, *τ*_*i*_ simply corresponds to the proportion of transcripts expressed by the *i*^th^ species which map to function, *T*. Assuming the community sequence spaces has been randomly sampled, these scenarios implicitly account for the number of gene copies (*t*) and the relative abundance (*p*) of the *i*^th^ species. In the case of metagenome-assembled genomes, relative contribution level can be calculated with equation 2 such that *t*_i_ is gene copies and abundance is measured as the relative proportion of reads mapping to single-copy marker genes for the *i*^th^ species.

As an example, imagine that we have a metagenome that has been binned into MAGs. Our hypothetical community has three species: a Cyanobacterium, Alphaproteobacterium, and a Gammaproteobacterium. Mapping metagenome reads to single-copy marker genes reveals that the Cyanobacteria, the Alphaproteobacterium and Gammaproteobacterium each represent 50%, 25%, and 25% of the community, respectively. We wish to calculate the functional redundancy of this community with respect to the production of a secondary metabolite. We use the presence of a gene, *xxxA*, as a proxy for an organisms’ potential to produce this secondary metabolite. The MAGs reveal that the Cyanobacterium and Alphaproteobacterium contain 5 copies of *xxxA* each, while the Gammaproteobacterium lacks the *xxxA* gene. Thus, the relative contribution from each species to secondary metabolite production, when using *xxxA* as a proxy, is *τ=*0.67, *τ=*0.33, and *τ*=0, respectively

Next, we reformulate the abundance distribution for community, *C*, as an abundance distribution with respect to *τ* (relative contribution distribution), such that:

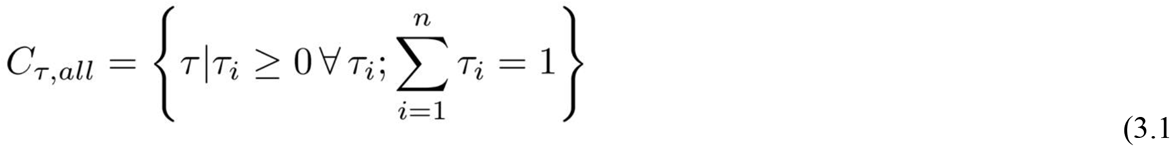

and

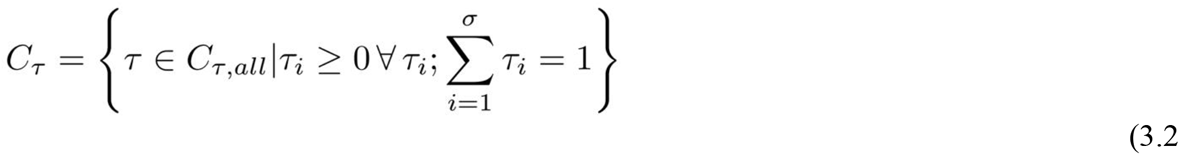

where again, *τ*_*i*_ is the relative contribution by the *i*^th^ species to *T*, such that all *τ* sum to equal 1. The community represented by equation 3.1 includes species not contributing to the community-aggregated parameter. The community represented by equation 3.2 is a subset of the one represented by equation 3.1, where *C*_*τ*_ contains only species from *C*_*τ,all*_ contributing towards a community-aggregated parameter, which we refer to as “contributors.” As such, we adopted the variable, *σ*, which represents the richness of contributors. The effective richness of contributors towards *T* was calculated using equation 3.2 and the Tsallis entropy index (21)(21) such that:

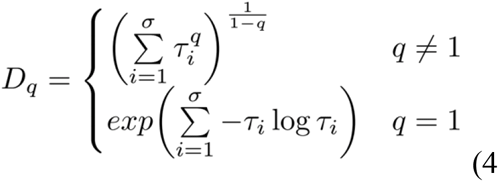

where *τ*_*i*_ is the relative contribution by *i*^th^ species in *C*_τ_ towards *T*, and *D*_q_ is the effective richness of contributors, and *q* is the diversity order, which weights the importance of species contributing more or less towards *T* when calculating *D*_q_. This is analogous to diversity order weighting the importance of rare versus abundant species when measuring diversity of species abundances (22). With diversity order *q*=0, equation 4 simplifies to the number of contributors in equation 3.2 (i.e., *σ*). Diversity order *q*=0 removes any influence in the variability of *τ* when calculating *D*_q_. Increasing the diversity order subsequently decreases the perceived contribution of contributors with smaller *τ*. Diversity orders *q*=1 and *q*=2 correspond to Shannon’s index. and Simpson’s index.

We then normalize Dq by community richness S, which is equal to Ds if every species were to contribute to T equally), and adjust evenness to account for the functional dependence that the lower evenness bound has on community richness (20)(20). As such, functional redundancy spans the full range from 0 to 1, such that:

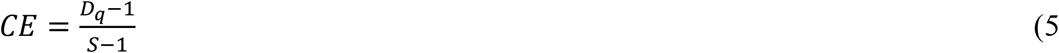

Here, functional redundancy, *CE*, indicates how evenly species contribute to a trait, *T*. For the hypothetical community described above, the effective richness (*D*_q_, *q*=0.5—see *Results*) of species possessing *xxxA* was ∼1.55, and the functional redundancy, with respect to *xxxA*, was *R*=0.47 (*q*=0.5; see *Results*).

### Simulations of Functional Redundancy as a Function of Trait Stability

In order to test the fidelity with which CE measures trait stability, or the stability of a community-aggregated parameter to species extinction, we simulated artificial communities with varying values of species richness (*S*), richness of contributors (*σ*), and contribution levels by contributors (*t)*. First, the richness of contributors was randomly chosen from a uniformly distributed range of 1 to 100. Next, the richness of species not contributing to the community-aggregated parameter was randomly chosen from a uniformly distributed range of 0 to 100, so that some artificial communities were dominated by contributors while others by non-contributors. The relative abundance, *p*, of individual species was modeled log-normally (28, 29)(28, 29) such that:

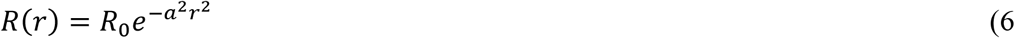

where *R* is the *r*^th^ rank, *R*_0_ is the maximum abundance, and *a* is the inverse width of the lognormal distribution. Values of *a* were randomly selected according to a uniform distribution from 0 (even) to 0.4 (more uneven), at 0.01 intervals. After establishing species relative abundances, we randomly chose *σ* contributing taxa from our lognormally distributed community. The contribution level, *t*, was then randomly assigned to these species from a uniformly distributed range of 0.01 to 1, at intervals of 0.01. We generated and calculated *R* for 1,000 artificial communities.

The stability of a community-aggregated parameter to extinction was tested by a simulation in which stability to extinction was quantified as the average fraction of species richness (*S*) that must go extinct in order to decrease community-aggregated parameter by 99%. Species were randomly removed from the community until summed *τ* was reduced to 1% of its original level. We used a 99% extinction, rather than 100%, to allow the evaluation of communities where all species were contributors. In such communities, all species would have to go extinct to remove 100% of the ecosystem function. We think that it is also justified ecologically, because dormant microbes in the rare biosphere can serve as a deep reservoir of traits (30), and it is not clear that fully removing a trait from an ecosystem happens in practice. The extinction simulation was performed 100 times for each of the individual 1,000 artificial communities. The stability to extinction generated from the 100 simulations were then averaged for each of the individual 1,000 artificial communities. We made no *a priori* assumptions about the optimal diversity order (i.e., *q*) to use in equations 4 and 5. We performed the simulation described above varying diversity orders from 0 to 2 at 0.1 intervals. The best diversity order was determined by fitting generalized additive model (GAM) regressions between *R* and stability to extinction, and then choosing the diversity order which resulted in a GAM fit with the lowest mean absolute error among residuals. All GAM fits were performed using the gam function from the R package mgcv (31). Last, we performed the same simulations and calculated *R* using other methods described in the literature. A summary of these measurements is provided in Table 1. An R package which calculates functional redundancy (equations 1-5) and performs these simulations is available at https://github.com/taylorroyalty/funfunfun.

**Table 1.** A summary of metrics of community-level functional redundancy.

| Authors | Interpretation | Equation | Notes |
| --- | --- | --- | --- |
| This work | Effective proportion of species with a trait | $R = \frac{D_q - 1}{S - 1}$ | • Accounts for species abundances<br>• Continuous trait states<br>• Individual traits<br>• $R$ spans 0 to 1 |
| (8) | Expected trait dissimilarity among species in a community | $R = 1 - \frac{\sum_i p_i \sum_j p_j \delta_{ji}}{D_2}$<br>$\delta_{ji}$ = trait distance<br>$D_2$ = taxonomic diversity order (q=2) | • Accounts for species abundances<br>• Continuous trait states<br>• Individual or multivariate traits<br>• Normalized by a maximum theoretical dissimilarity<br>• $R$ spans 0 to 1 |
| (6) | Ratio of species to number of functional entities (states) | $R = \frac{S}{FE};$<br>FE = # of possible combinations of trait states for all traits under consideration | • Binary trait states<br>• Individual or multivariate traits<br>• $R$ spans $\sim 0 (\frac{S}{FE} = 0)$ to $S$ |
| Various (see text for examples) | Fraction of species with a trait | $R = \frac{v}{S}$ | • Binary trait states<br>• Individual traits<br>• $R$ spans 0 to 1 |
| (7) | Median overlap of traits between species | $R = med(O);$<br>$O = \{o_{i,j} = \vert g_i \cap g_j\vert ; i \neq j\};$<br>$g = taxa;$<br>$O$ = set containing # of overlaps among species | • Binary trait states<br>• $\vert g_i \cap g_j\vert $ represents the number of overlapping traits $g$ between species $i$ and species $j$<br>• Individual and multivariate traits<br>• $R$ spans 0 to ∞ |

### A Case Study of Functional Redundancy in the Global Ocean Nitrogen Cycle

Draft metagenome-assembled genomes (MAGs) from Tully, Graham et al. (24) were downloaded from NCBI (32). Data on metagenome sequencing effort (number of short reads), MAG relative abundance estimates (pre-corrected for genome size), ocean depths, and ocean regions, defined in (24), were downloaded from Figshare (https://doi.org/10.6084/m9.figshare.5188273). MAG relative abundances from each site were multiplied by the respective site’s sequencing effort to convert relative abundances into abundance count data. Abundance counts for all sites were normalized by the lowest abundance (greater than 0) among all sites. MAG abundances were then rarefied using the rrarefy function from the R package vegan (33). We chose a sampling effort of 14,265, as this sampling effort corresponded to the site with lowest sampling effort (i.e., sequencing effort). We tabulated nitrogen-cycle marker genes in individual MAGs by blasting MAG proteins against nitrogen-cycle marker gene databases using the BLASTP algorithm (*E*-value=10^−10^) (34). Nitrogen cycle marker genes were categorized into eight categories: N_2_ fixation (*nifH*), nitrification (*amoA, hao*), NO_3_ reduction (*napA, narG*), dissimilatory NO_3_ reduction (*nrfA*), assimilatory NO_3_/NO_2_ reduction (*narB, nasA, nasB*, and *nirB*), NH_4_^+^ assimilation (*gdhA, glnA*), N remineralization (*gltB*), and denitrification (*nirK, nirS, norB*, and *nosZ*) (23). Individual BLAST databases were constructed for each nitrogen cycle category. Gene names associated with individual categories were searched for in the Swiss-Prot database (35). All proteins matching gene names were subsequently downloaded and used in BLAST database construction. Functional redundancy was then calculated for individual stations, at individual depths, for each nitrogen cycle category using equations 1-5. We used a diversity order, *q*=0.5, based on the GAM sensitivity analysis. Table 2 contains protein names, gene names, corresponding EC, and entry name for all genes used in the above analysis.

**Table 2.**
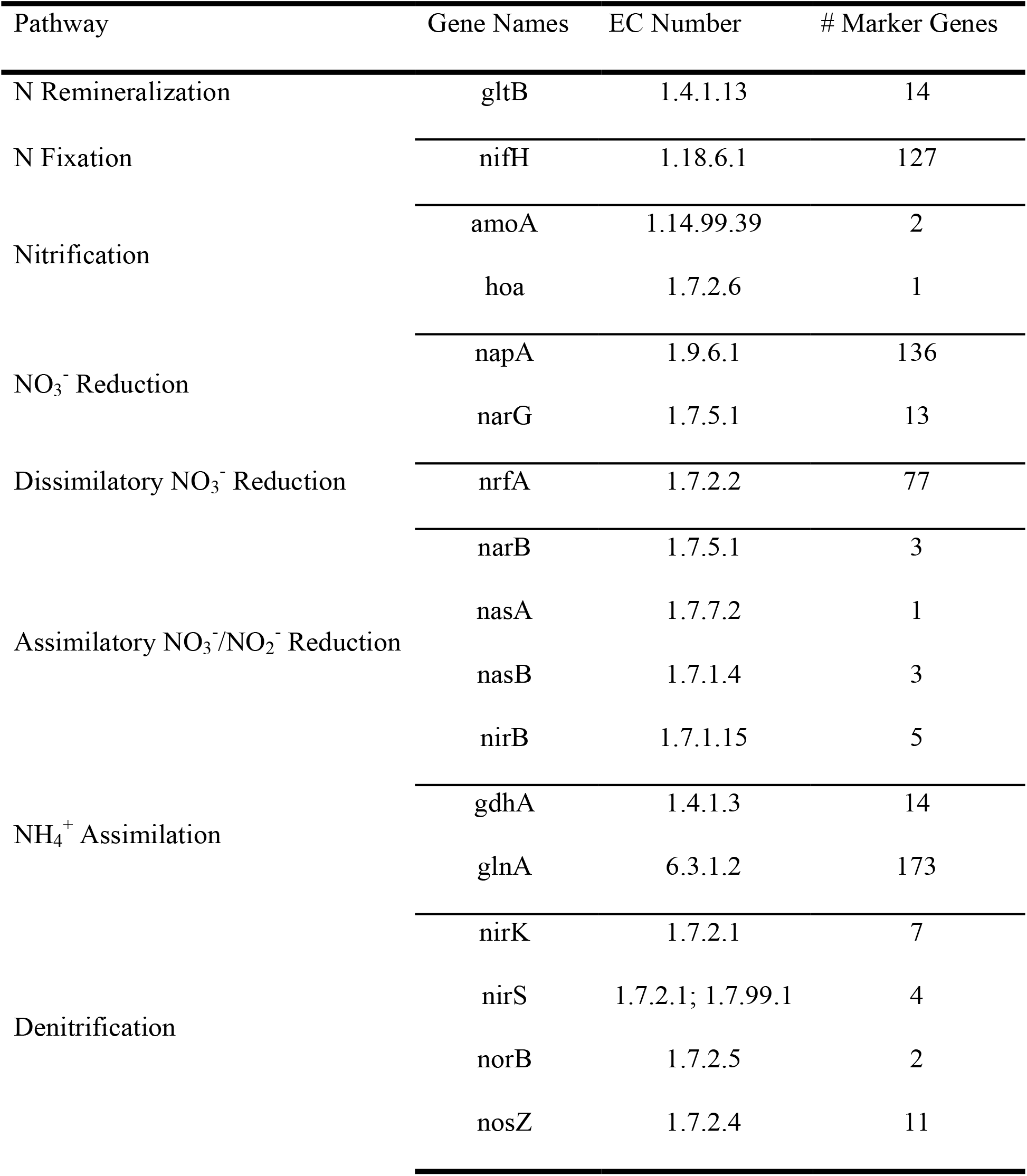
A summary of marker gene sequences used in the BLASTP analysis of *TARA Oceans* MAGs. All marker genes were downloaded from Swiss-Prot.

## Results and Discussion

### Representing microbial communities as species abundance, trait abundance, and relative contribution distributions

CE represents functional redundancy using two distributions: the distribution of species abundances within a community, and the distribution of contribution levels among species. Here, we evaluate genomic traits because these are regularly adopted as proxies for microbially-mediated biogeochemistry (16, 26, 36–38). Because trait levels are measured as non-negative, real numbers (18), the relative contribution of individual species to a community-aggregated parameter can be represented as a relative abundance distribution (Figure 1). We determined the relative contribution of individual species to the community-aggregated parameter from the product of species relative abundances and their respective trait abundance distributions (equation 2). We refer to the distribution of the relative contribution of each species to total as the relative contribution distribution (equation 3). Thus, an abundant species that expresses a small amount of trait per cell, and a rarer species that expresses a large amount of trait per cell, can contribute equally with respect to a community-aggregated parameter (Figure 1).

**Figure 1:**
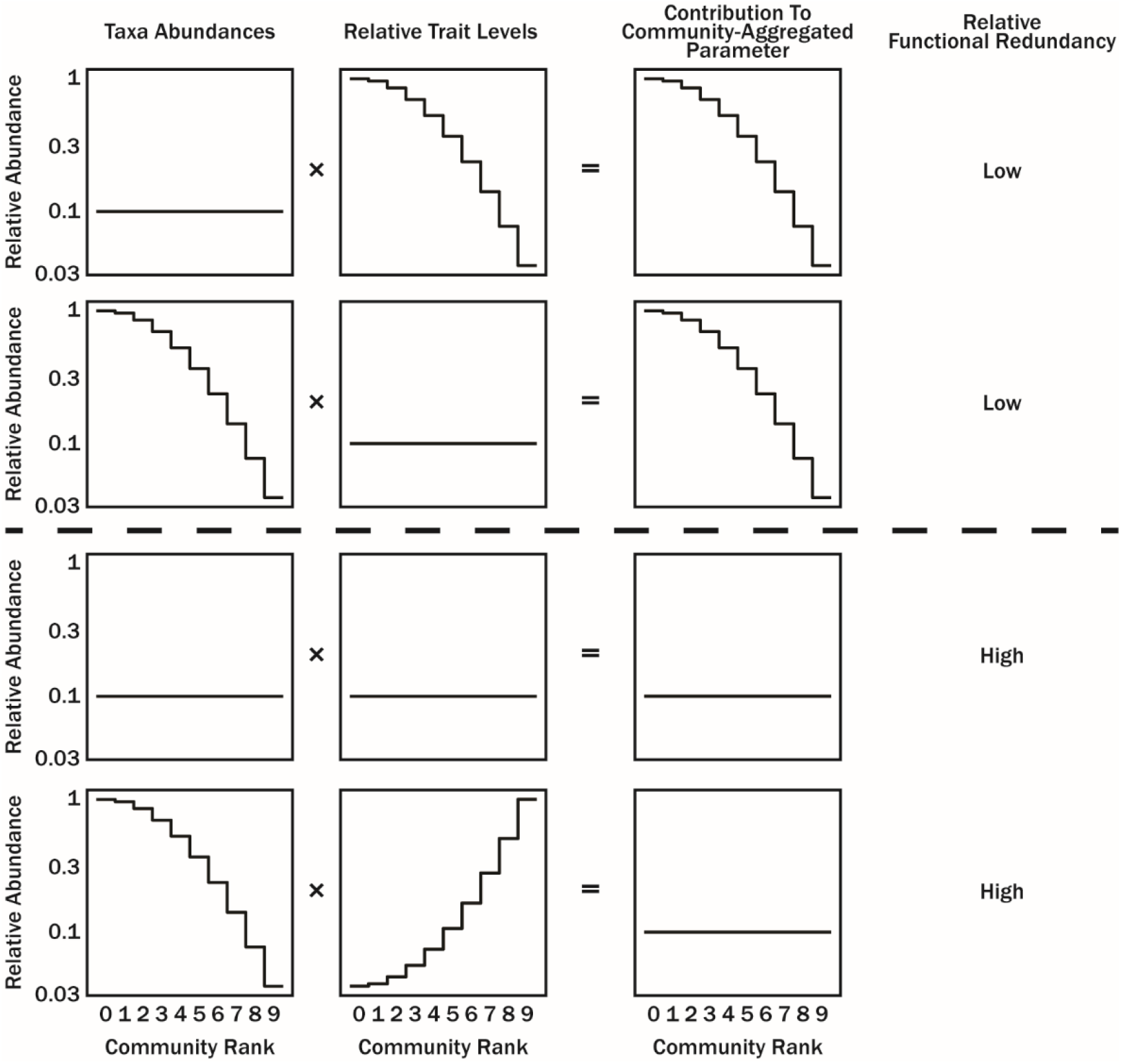
Community taxon abundance and relative contribution level are scalable and provide a measure for the contribution that different species have towards a community-aggregated parameter. The evenness of species-level contribution to the community-aggregated parameter is taken as the functional redundancy.

### Defining, Quantifying Functional Redundancy, and Choosing a Diversity Order q

We measured functional redundancy as how evenly species in a community contribute to a community-aggregated parameter and termed this metric as contribution evenness (CE). This definition is consistent with descriptions of functional redundancy as multiple species sharing similar ecological roles or functions (2, 3). The relative contribution by species towards a community aggregated-parameter (equation 2) provides a framework to calculate the effective number, or richness, of species that contribute to that community-aggregated parameter as an ecological function. This effective richness is calculated with traditional diversity indices (equations 3-4) (21, 22). To determine the effective proportion of a community contributing to the community-aggregated parameter, the richness of contributors is normalized by species richness (22). Last, to make meaningful comparisons of evenness between communities with different levels of richness, we made our evenness metric unrelated to community richness (equation 5)(20).

Numerous diversity indices have been proposed in the literature. Three common measures of diversity include species richness, Shannon’s diversity index, and Simpson’s diversity index, are manifestations of equation 4 when the *q* term—commonly referred to as the diversity order—is *q*=0, *q*=1, and *q*=2, respectively. The diversity order changes the relative weight placed on rare versus common species when determining the effective richness. We found that GAM fits between functional redundancy and stability to extinction followed positive, monotonic functions for all diversity orders spanning *q*=0 to *q*=2. However, at *q* ≈ 0.5, CE was most tightly related to trait stability, our benchmark for functional redundancy efficacy (Figure 2; also compare Fig 3a to Fig 3b). Therefore, we suggest that CE is most effective as an indicator of trait stability *q*=0.5 makes *CE* the best indicator of how resilient traits are to species extinction. In that case, equation 4 simplifies to:

**Figure 2:**
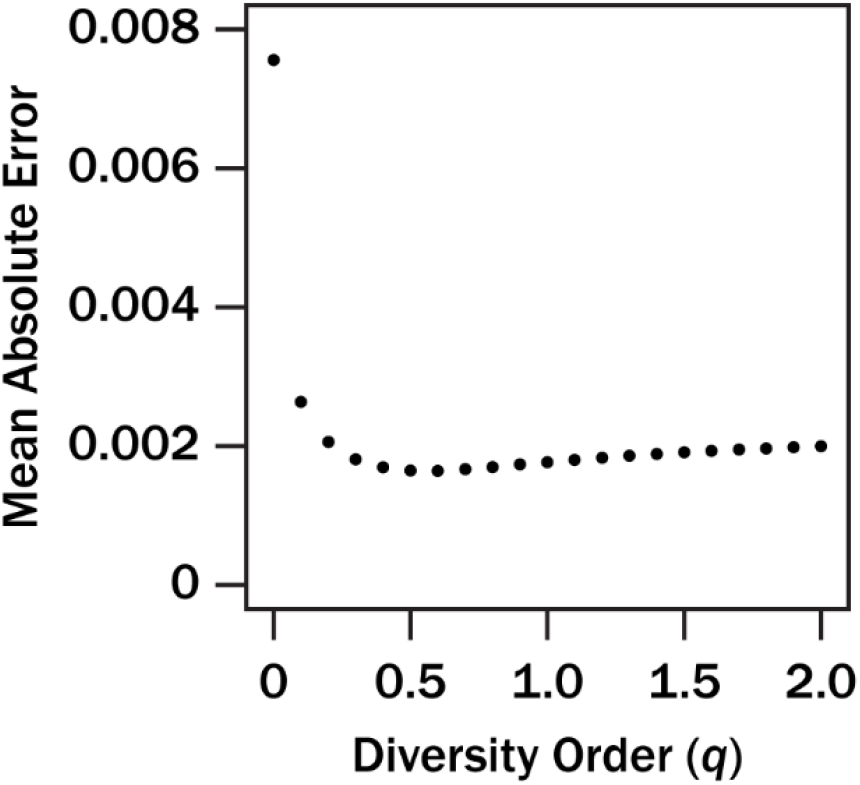
The sensitivity of mean absolute error between functional redundancy and trait stability to species extinction in response changing diversity order *q* in equation 4.

**Figure 3:**
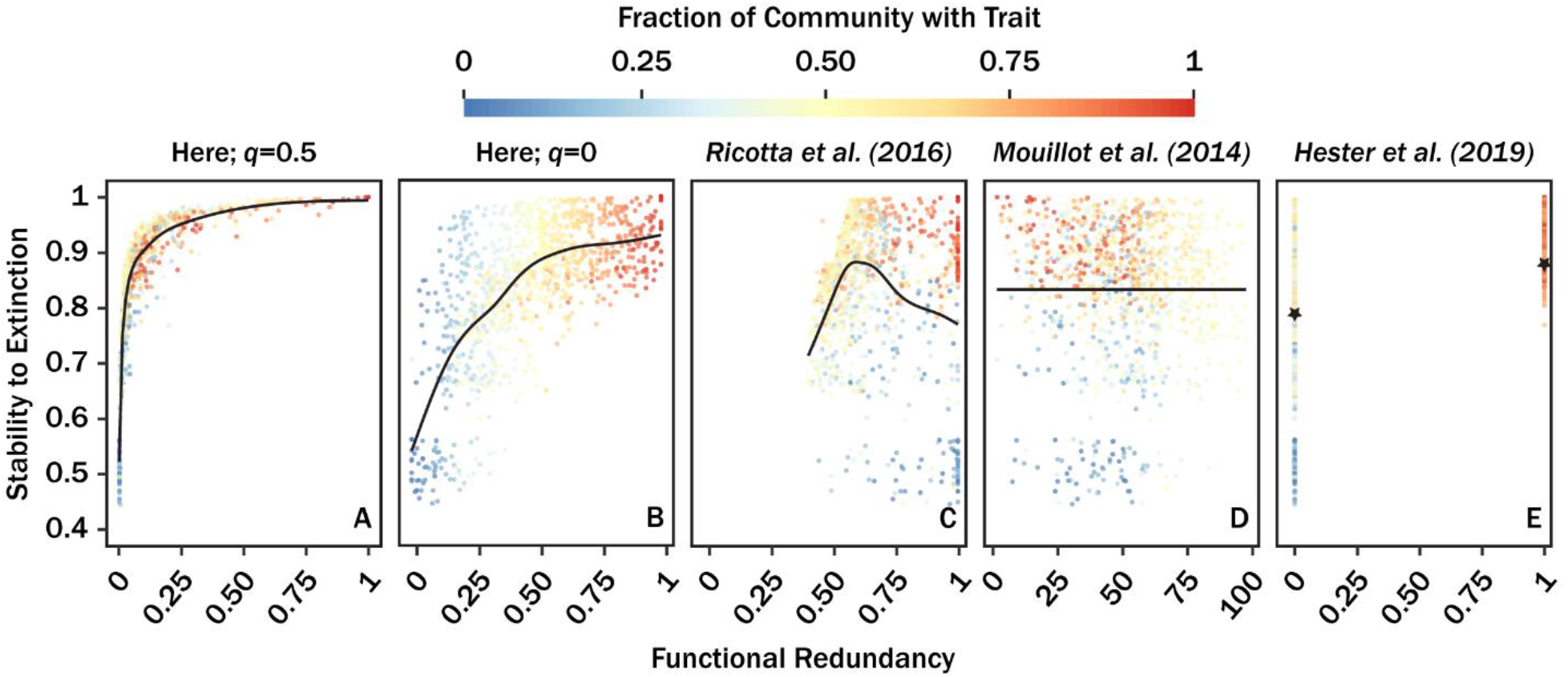
A comparison of the relationship between selected different functional redundancy metrics and trait stability (see methods). Functional redundancy calculated with equation 5 (*q*=0.5) (**A**), equation 5 (*q*=0) (**B**), methodology from Ricotta et al. (2016) (**C**), methodology from Mouillot et al. (2014) (**D**), and methodology from Hester et al. (2019). Black lines in panels **A-D** correspond to best fit GAMs. The stars in panel **E** correspond to the average trait stability at *R*=0 and *R*=1. The color scale corresponds to the fraction of a community with a trait versus.

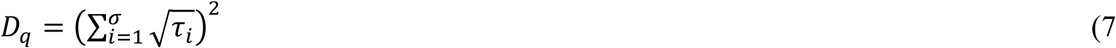

and CE becomes:

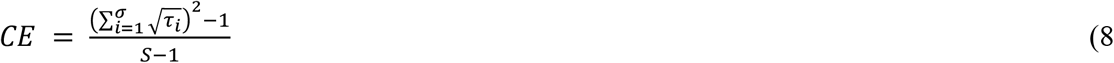

This is the formulation of CE that we recommend for general use.

### Functional Redundancy and Trait Stability to Species Extinction

A tangible ecological consequence of functional redundancy is stability of an ecosystem function to the extinction of species performing that function. When functions are more broadly distributed across species, a function is less likely to disappear due to the extinction of any subset of species. This interpretation is often highlighted in macro-ecological studies (2–4, 7, 39–41). The conceptual relationship between functional redundancy and stability of an ecosystem function provides an opportunity to demonstrate the fidelity of CE. To do this, CE was calculated for randomly generated communities. Individual species were randomly assigned relative contribution levels towards a community-aggregated parameter. We simulated random species extinction events in the artificial communities until >99% of the original ecosystem function was removed from the community (see *Methods*).

It is important to emphasize that these simulations were not designed to mimic how extinction occurs in nature. Instead, these simulations empirically demonstrate the degree to which an ecosystem function is concentrated in species. The more widely an ecosystem function is distributed across species, the more species must go extinct before the ecosystem function is lost. We observed a positive, monotonic relationship between the stability to random species extinction events and CE (Figure 3A). When CE was ∼0, stability to species extinction was ∼0.5, meaning ∼50% of the community, on average, went extinct before an ecosystem function was removed from the community. This is equivalent to only one species performing the ecosystem function. When CE was ∼1, stability to species extinct was ∼1, meaning that 100% of the community, on average, had to go extinct to completely remove a function from an ecosystem. In general, the relationship between CE and trait stability to extinction resembled a saturation curve—although we opted to use GAMs when calculating mean absolute error (previous section) as we made no assumption about the true relationship between CE and function stability, other than it should be a positive, monotonic function.

Here, we recommend the form of CE described in equation 8, as it is optimized for our intended use case, which is measuring relative stability of traits to species extinctions within communities. However, we can imagine other applications of the CE concept, for which other values of *q* may be appropriate. Even for species diversity (i.e., equation 4), there is still discussion over the merits of using Simpson’s Index (*q*=2), Shannon’s Index (*q*=1), and richness (*q*=0) in different situations (42–44). As such, it is important to be explicit about the implications of behind choosing a trait and the diversity order.

### CE Measures Trait Stability More Reliably than Other Functional Redundancy Metrics

Several methods for evaluating community-level functional redundancy exist in the literature, as detailed in Table 1. While Table 1 certainly does not include all published metrics, the highlighted metrics reflect the diversity in how traits, community properties (i.e., richness and evenness), as well as the desired metric utility (by design) vary among different functional redundancy metrics. Two of the metrics evaluate niche overlap, where niches are defined in multivariate trait space (6, 7). In the case of Ricotta et al. (7), functional redundancy weights species abundance, leading to a measure of the expected niche dissimilarity from the mean community niche. In contrast, Hester et al. (6) evaluates the median metabolic overlap which considers neither species abundance nor trait levels. One metric evaluates niche occupancy, where niches are defined in multivariate trait space (5). In this approach, functional redundancy is taken to be as the ratio of species richness to plausible number niches (unique combinations of trait states), called functional entities. The last metric is informal but has been utilized by various authors. In this approach, functional redundancy is simply interpreted as the proportion of species in a community with a trait (e.g. Wohl et al., 2004; Tully et al., 2017; Trivedi et al., 2020).

To highlight the difference between our metric and the others described in Table 1, we repeated the extinction simulations and compared all estimates of functional redundancy with trait stability to species extinction (Figure 3B-E). In these simulations, only one trait was considered when calculating functional redundancy with all metrics. For the method presented in Mouillot et al. (5), we assumed there were two functional entities (trait absence and presence). Generally, all functional redundancy metrics trended differently to stability to species extinction. This observation was not surprising, as most of these methods evaluate niche overlap in some fashion. Among the metrics in Table 1, the fraction of the community possessing a trait behaved most like CE (Figure 3B). We note that the fraction of the community possessing a trait is a special case of CE, with diversity order *q*=0. Lastly, one inherent limitation of CE is that it only evaluates single community-aggregated parameters at a time. To address functional redundancy with respect to multiple traits perform dimensional reduction (e.g., PCA) on multiple traits and use the resultant factors as trait levels in equation 2. Factors in this case would be interpreted as different niches.

### Case Study: Redundancy of Nitrogen Cycle Marker Genes in Marine Microbiomes

To demonstrate a real application and provide a positive control for CE results, we measured CE in the marine nitrogen cycle using genomic data from the *TARA Oceans* project (47)(45). Recently, (24) published 2,361 draft metagenome-assembled genomes (MAGs) which represent genomic content from seven oceanographic provinces, three different depths, and four filter size fractions. Relative abundance data used here were reported in (24) and were determined by the proportion of sequence reads mapping to assembled MAGs. For simplicity, we limited our analysis to only MAGs identified in the bacterial size fraction (<3 *μ*m), which corresponds to free-living microbes. The nitrogen cycle was simplified to eight pathways (23)(23) and included N_2_ fixation, nitrification, NO_3_^-^ reduction, dissimilatory NO_3_^-^ reduction, assimilatory NO_3_^-^/NO_2_^-^ reduction, NH_4_^+^ assimilation, N remineralization, and denitrification. A general summary of the marker genes (downloaded from Swiss-Prot; see Methods) used for quantifying nitrogen cycle pathways in MAGs is summarized in Table 2.

Overall, the functional redundancy for individual genomic traits was consistent with our conceptual understanding of primary nitrogen-transforming pathways in the ocean (Figure 4). A clear example was the low functional redundancy in the nitrification pathway. We expected this observation as nitrification metabolisms in oceans are primarily restricted to *Crenarchaeota* (46). Similarly, the N_2_ fixation pathway, which is predominantly facilitated by a small group of *Cyanobacteria*, exhibits low functional redundancy. Conversely, the uptake of inorganic N via NH_4_^+^ and NO_3_^-^/NO_2_^-^ assimilation are known to be broadly expressed traits, and accordingly exhibit high CE. Last, the low functional redundancy in dissimilatory NO_3_^-^ reduction and NO_3_^-^ reduction were not surprising considering these pathways occur in anaerobic and low-oxygen environments, respectively (23).

**Figure 4:**
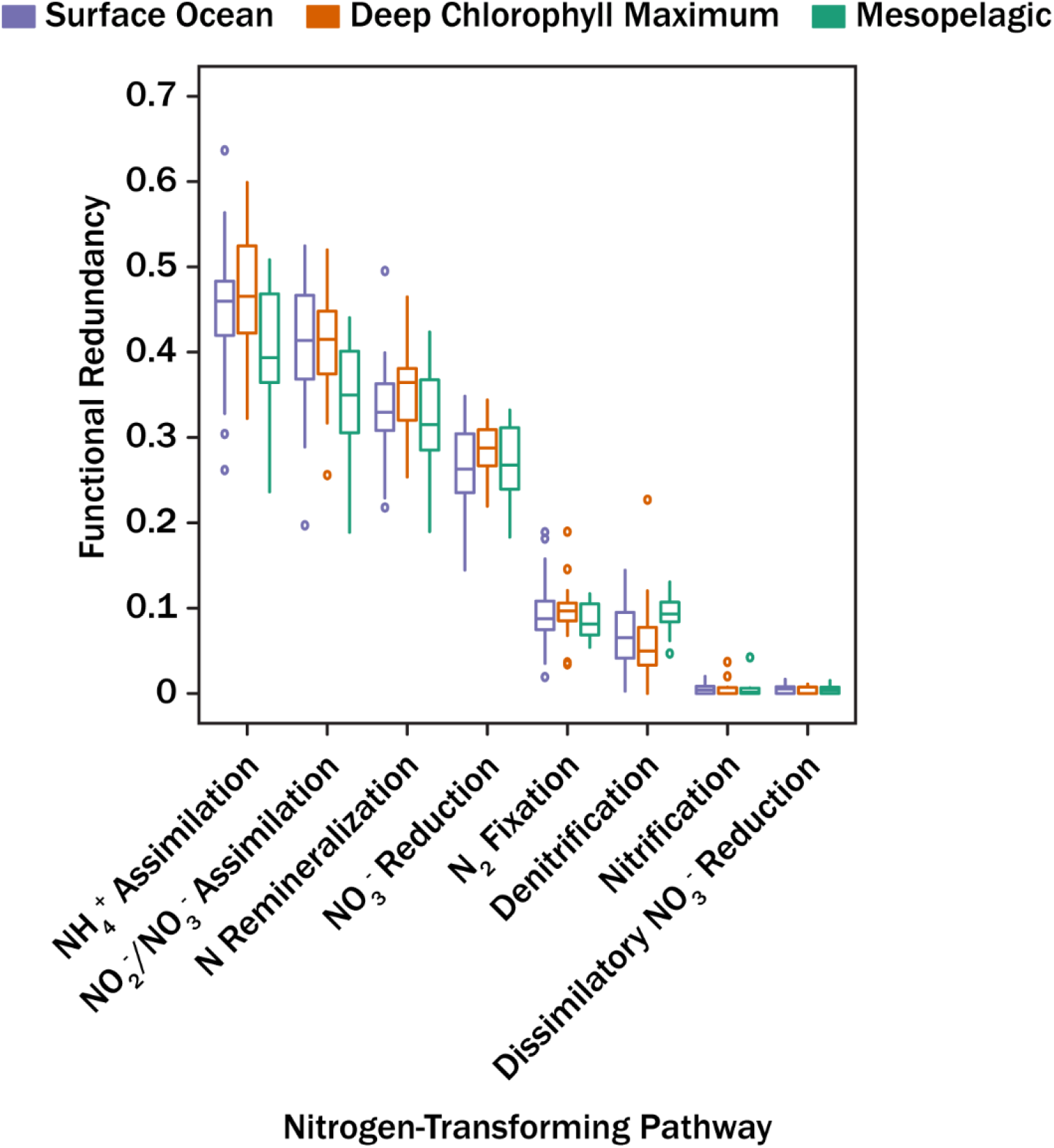
Boxplots showing functional redundancy calculated for eight nitrogen-transforming pathways in *TARA Oceans* draft MAGs. Different colors correspond to sample depth. Whiskers correspond to 150% of the inter-quantile region.

Some variance existed across oceanographic provinces and depth for intra-pathway functional redundancy. However, inter-pathway variance in functional redundancy was generally larger than intra-pathway variance (Figure 4). This suggests that the marine nitrogen cycle marker genes considered here are consistently distributed among communities for MAGs retrieved in Earth’s ocean at middle and lower latitudes. This was consistent with the lack of relationship between the metagenome companion metadata (e.g., O_2_, inorganic N, PO_4_^3-^, latitude, longitude, etc.) and functional redundancy. The low intra-pathway variance in functional redundancy was not particularly surprising, as saltwater biomes exhibit the lowest alpha-diversity and small beta-diversity among free-living microbiomes (47, 49)(45)(47). Minor shifts in community composition among lower taxonomic ranks (e.g., genus or species) will not drastically alter redundancy in genomic traits as changes in genomic composition is buffered by traits conserved at higher taxonomic ranks (e.g., phylum level) (50)(48). This observation was in agreement with an earlier analysis evaluating cluster of orthologous groups (COGs) composition across *TARA Oceans* metagenomes (45).

The disproportionate number of lineages that lack cultured representatives highlights the continued importance of nucleotide sequencing in microbial ecology and the interference of microbial function (49). Here, we demonstrated that CE provides a framework to evaluate the functional redundancy for these traits in a novel way. This approach represents an easy-to-apply, quantitative metric for functional redundancy that faithfully represents the stability of traits to extinctions of species within a community.

## Data Availability

In order to facilitate the use of CE functional redundancy measurements, we created an R package to calculate CE functional redundancy and run extinction simulations. The package is available at https://github.com/taylorroyalty/funfunfun.

## Acknowledgments

We gratefully acknowledge the comments of three anonymous reviewers, whose constructive comments substantially improved the manuscript. Funding for this project was provided by a C-DEBI graduate fellowship to TMR and by the U.S. Department of Energy, Office of Science, Office of Biological and Environmental Research, Genomic Science Program under Award Number DE-SC0020369 for TMR and ADS. This is C-DEBI contribution number [to be assigned upon manuscript acceptance].

## Author Contribution

TMR designed the research, performed research, analyzed data, developed the R package, and wrote the paper. ADS designed the research, developed the R package, and wrote the paper.

